# Multi-Scale Network Regression for Brain-Phenotype Associations

**DOI:** 10.1101/628651

**Authors:** Cedric Huchuan Xia, Zongming Ma, Zaixu Cui, Danilo Bzdok, Danielle S. Bassett, Theodore D. Satterthwaite, Russell T. Shinohara, Daniela M. Witten

## Abstract

Complex brain networks are increasingly characterized at different scales, including global summary statistics, community connectivity, and individual edges. While research relating brain networks to demographic and behavioral measurements has yielded many insights into brain-phenotype relationships, common analytical approaches only consider network information at a single scale, thus failing to incorporate rich information present at other scales. Here, we designed, implemented, and deployed Multi-Scale Network Regression (MSNR), a penalized multivariate approach for modeling brain networks that explicitly respects both edge- and community-level information by assuming a low rank and sparse structure, both encouraging less complex and more interpretable modeling. Capitalizing on a large neuroimaging cohort (*n* = 1, 051), we demonstrate that MSNR recapitulates interpretable and statistically significant association between functional connectivity patterns with brain development, sex differences, and motion-related artifacts. Notably, compared to single-scale methods, MSNR achieves a balance between prediction performance and model complexity, with improved interpretability. Together, by jointly exploiting both edge- and community-level information, MSNR has the potential to yield novel insights into brain-behavior relationships.

## 1. Introduction

Studying brain-phenotype relationships in high-dimensional connectomics is an active area of research in neuroscience (Bassett and Sporns 2017; Bzdok et al. 2016). The advent of large neuroimaging datasets that provide measures of brain connectivity for unprecedented numbers of subjects have yielded novel insights into development and aging, cognition, and neuropsychiatric illnesses (Biswal et al. 2010; Van Essen et al. 2012; Bzdok and Yeo 2017; Schumann et al. 2010; Jernigan et al. 2016). As the availability of datasets with rich neural, genetic, and behavioral measurements from large numbers of subjects continues to increase, there is a growing need for statistical methods that are tailored for the discovery of complex relationships between brain networks and phenotypes (Craddock, Tungaraza, and Milham 2015; Varoquaux and Craddock 2013).

A typical brain network consists of hundreds of nodes that denote brain regions, and tens of thousands of edges that indicate connections between pairs of nodes (Rubinov and Sporns 2009). The network can be viewed on the *micro*-scale, *meso*-scale, or *macro*-scale. The micro-scale of the network can be characterized by features of its edges. The *macro* - scale of the network can be characterized by global features such as characteristic path length and global efficiency (Rubinov and Sporns 2009). The *meso*-scale falls in between the microscale and macro-scale, and includes the communities that make up the network (Sporns and Betzel 2016; Betzel, Medaglia, and Bassett 2018). A community refers to a collection of nodes that are highly connected to each other and have little connection to nodes outside of the community. Prior work has demonstrated that brain network architecture present on these different scales is associated with development and aging (Power et al. 2010; Gu et al. 2015; Betzel et al. 2014), cognition (Crossley et al. 2013; Park and Friston 2013; Bressler and Menon 2010), and neuropsychiatric diseases (Yu et al. 2019; Braun et al. 2016; Fornito, Zalesky, and Breakspear 2015; Grillon et al. 2013; Bassett, Xia, and Satterthwaite 2018; Xia et al. 2018; Kernbach et al. 2018).

Despite increasing appreciation that multi-scale organization of the brain may be responsible for some of its major functions (Betzel and Bassett 2017; Bassett and Siebenhühner 2013), thus far, common strategies for studying the relationship between brain connectivity and phenotypes tend to consider network features at a single scale (Craddock, Tungaraza, and Milham 2015; Varoquaux and Craddock 2013). For example, a popular single-scale strategy focuses on group-level comparisons of individual connections in brain networks (Grillon et al. 2013; Fornito, Zalesky, and Breakspear 2015; Bressler and Menon 2010). This approach typically involves performing a statistical test on each network edge. While this procedure is easy to implement, several drawbacks limit its effectiveness (Bzdok and Ioannidis 2019). Two main limitations are the need to account for multiple comparisons, and a lack of interpretability (Craddock, Tungaraza, and Milham 2015; Varoquaux and Craddock 2013). To achieve high power while minimizing the risk of false positives, alternative edge-based methods have been developed, such as the network-based statistic (Zalesky, Fornito, and Bullmore 2010) and multivariate distance matrix regression (Zapala and Schork 2012). While these strategies have yielded important insights, they nonetheless focus exclusively on the micro-scale, often producing results that are difficult to interpret and that do not exploit the multi-scale information present in the brain networks.

Given the importance of community structure in brain networks and its interpretability in the context of neural and cognitive computations (Sporns and Betzel 2016; Betzel, Medaglia, and Bassett 2018), it might be tempting to conduct a mass-univariate analysis at the meso-scale, considering *within* - and *between*-community connectivity as the input features (Yu et al. 2019; Betzel et al. 2014; Gu et al. 2015; Braun et al. 2016). Such an approach dramatically reduces the dimensionality of the data, which in turn decreases the burden of multiple comparisons correction. A community-based approach also has the added benefit of not having to deconstruct the connectivity matrix into vectors, as in an edge-based approach, which inevitably disrupts the innate data structures. However, summarizing thousands of edges as one single number to represent the connection within or between communities can be problematic, especially for large communities such as the default mode network, whose edges are spatially distributed across the anterior and posterior portions of the brain as well as across both medial and lateral surfaces (Raichle 2015). Stated differently, extracting the mean connectivity at the community level risks mixing disparate signals.

In this paper, we introduce *Multi-Scale Network Regression (MSNR)*, which simultaneously incorporates information across multiple scales in order to reveal associations between high-dimensional connectomic data and phenotypes of interest. We first describe the MSNR model and introduce an algorithm to estimate the parameters. Next, we capitalize on one of the largest neurodevelopmental imaging cohorts, the Philadelphia Neurodevelopmental Cohort (PNC), to empirically assess the ability of MSNR in delineating brain connectivity patterns that are associated with a wide variety of phenotypes. Importantly, we conduct head-to-head comparisons between MSNR and common edge- and community-based analyses that are based on single-scale strategies, and show that MSNR achieves a balance between prediction performance and interpretability by considering information at multiple network scales.

## 2. Statistical Methodology

### 2.1. A Statistical Model for Multi-Scale Network Regression

Given *n* subjects, let *A*^1^,…,*A^n^* ∈ ℝ^*p×p*^ denote the adjacency matrices corresponding to their brain networks, where *p* is the number of nodes. For instance, 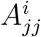 could represent the Pearson correlation of the mean activation timeseries of two brain regions, a common measure of functional connectivity, between the *j*-th and *j*′-th nodes for the *i*-th subject. Furthermore, we assume that the *p* nodes can be partitioned into *K* distinct communities *C*_1_,…,*C_K_* that are known *a priori*: 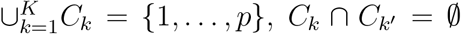 if *k* ≠ *k*′. The notation *j* ∈ *C_k_* indicates that the *j*-th node is in the *k*-th community. Moreover, for each subject, *q* covariates have been measured, so that 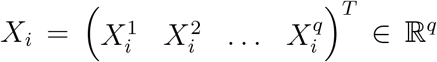 is a covariate vector for the *i*-th subject, *i* = 1,…, *n*.

In what follows, we consider the model

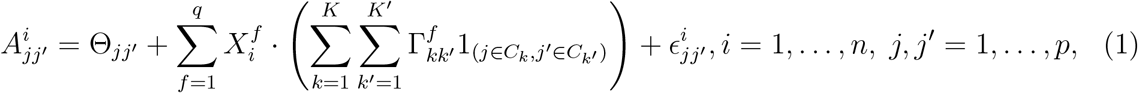

where 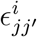 is a mean-zero noise term, and 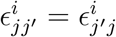. The matrix Θ is a symmetric *p×p* matrix that summarizes the mean connectivity (across all of the subjects) of each pair of nodes, in the absence of covariates. Finally, for *f* = 1,…,*q*, Γ^*f*^ is a symmetric *K × K* matrix that quantifies the association between the *f*-th feature and the connectivity between each pair of communities. For instance, a one-unit increase in 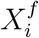 is associated with a 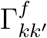 increase in the mean connectivity between nodes in the *k*-th and *k*′-th communities.

We now define a *p × K* matrix *W* for which *W_jk_* = 1_(*j∈C_k_*_), where 1_(·)_ denotes an indicator variable. As such, (1) can be re-written as

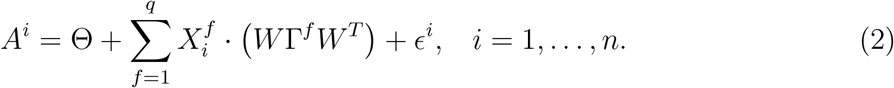

In order to fit the model (2), we make two assumptions about the structures of the unknown parameter matrices Θ and Γ^1^,…, Γ^*q*^.

#### Assumption 1

Θ *has low rank* (Smith et al. 2015; Leonardi et al. 2013; Li et al. 2009). That is, Θ = *VV^T^* where *V* is a *p × d* matrix, for a small positive constant *d*. This means that the *p* nodes effectively reside in a reduced subspace of *d* dimensions. The mean connectivity between any pair of nodes is simply given by their inner product in this low-dimensional subspace.

#### Assumption 2

Γ^1^,…,Γ^*q*^ *are sparse* (Meunier, Lambiotte, and Bullmore 2010; Newman 2006; Xia et al. 2018). That is, most of their elements are *exactly* equal to zero. If 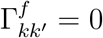, then the value of the *f*-th feature is unassociated with the mean connectivity between nodes in the *k*-th and *k*′-th communities.

We note that Assumption 1 is closely related to the random dot product graph model and similar models (Fosdick and Hoff 2015; Durante, Dunson, and Vogelstein 2017; Durante and Dunson 2018; Tang et al. 2017; Young and Scheinerman 2007), whereas Assumption 2 is a standard sparsity assumption for high-dimensional regression (Tibshirani 1996; Hastie et al. 2015; Hastie, Tibshirani, and Friedman 2008). Under these two assumptions, a schematic of the model (2) can be seen in Fig. 1.

**Figure 1:**
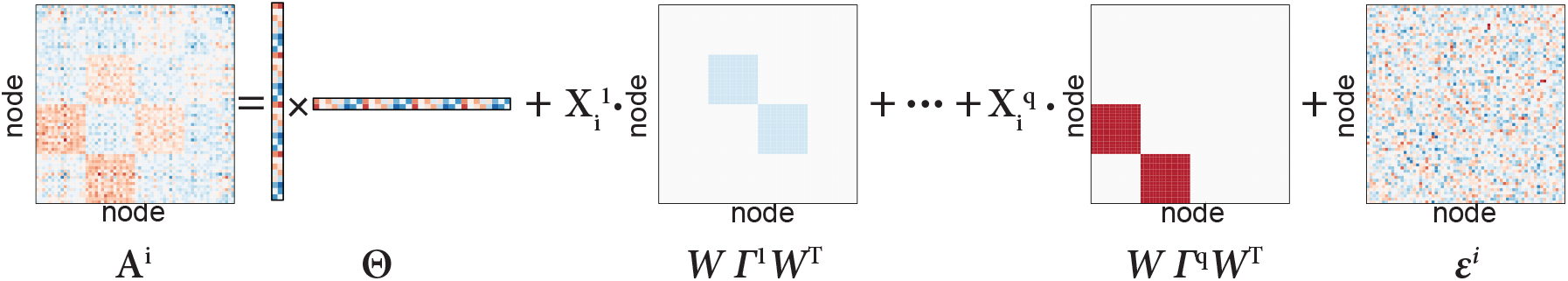
A schematic for Multi-Scale Network Regression. Under model (2), *A^i^* is the adjacency matrix for the *i*-th subject, Θ is a low-rank matrix representing the mean connectivity across all subjects, Γ^1^,…,Γ^*q*^ are sparse matrices representing the community-level connectivity associated with the covariates 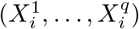, and *ϵ^i^* is the noise.

Model (2) is closely related to both the stochastic block model (Choi, Wolfe, and Airoldi 2012) and the random dot product graph model (Young and Scheinerman 2007). In particular, if Θ = 0, *q* =1, and 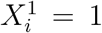 for *i* = 1,…, *n*, then (2) reduces to a stochastic block model with known communities *C*_1_,…, *C_K_*. And if Γ^1^ = … = Γ^*q*^ = 0 and Assumption 1 holds, then (2) reduces to a random dot product graph model. However, unlike those two models, model (2) explicitly allows for the mean of the adjacency matrix to be a function of covariates, and effectively incorporates both edge- and community-level network information.

### 2.2. Optimization Problem

We now consider the task of fitting model (2), under Assumptions 1 and 2. It is natural to consider the optimization problem

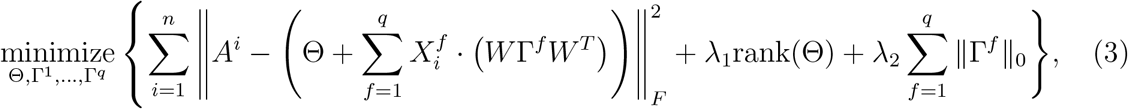

where the notation 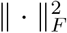 indicates the squared *Frobenius* norm of a matrix, i.e. 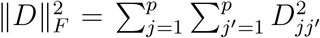, and the notation ||·||_0_ indicates the element-wise cardinality (or *ℓ*_0_ norm) of a matrix, i.e. 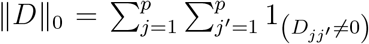. In (3), λ_1_ and λ_2_ are non-negative tuning parameter values that control the rank of Θ and the sparsity of Γ^1^,…, Γ^*q*^, respectively.

Unfortunately, due to the presence of the rank and *ℓ*_0_ penalties, the optimization problem (3) is highly non-convex, and no efficient algorithms are available to solve it. Thus, in what follows, we will consider an alternative to (3), which results from replacing the non-convex rank and *ℓ*_0_ penalties in (3) with their convex relaxations. This reformulation of the objective function leads to the optimization problem

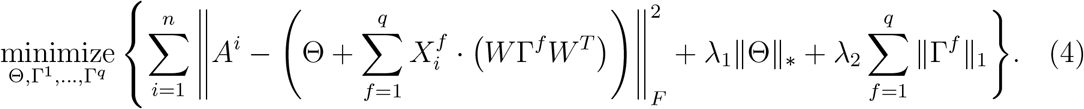

In (4), the notation ||·||_*_ indicates the *nuclear norm* of a matrix, i.e. the sum of its singular values (Fazel 2002; Recht, Fazel, and Parrilo 2010; Bien and Witten 2016). The nuclear norm is a convex surrogate for the rank of a matrix. The notation ||·|_1_ indicates the element-wise *ℓ*_1_ norm of a matrix, i.e. 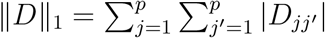; this is a convex relaxation of the *ℓ*_0_ norm (Tibshirani 1996; Hastie et al. 2015; Hastie, Tibshirani, and Friedman 2008). In (4), the non-negative tuning parameters λ_1_ and λ_2_ encourage Θ and Γ^1^,…,Γ^*q*^ to be low-rank and sparse, respectively.

Importantly, the optimization problem (4) is convex, and so fast algorithms are available to solve it for the global optimum. In Section 2.3, we derive a block coordinate descent algorithm for solving (4).

### 2.3. Block Coordinate Descent Algorithm to Solve (4)

We now derive a block coordinate descent algorithm for solving (4) (Tseng 2001; Hastie, Tibshirani, and Friedman 2008; Bien and Witten 2016; Friedman et al. 2007). Roughly speaking, we will cycle through the parameters Θ and Γ^1^,…, Γ^*q*^, and minimize the objective (4) with respect to each one in turn, holding all others fixed. Because the loss function is differentiable and the penalties are separable with respect to each block of parameters, this approach is guaranteed to reach the global optimum. The algorithm is as follows:

1. Initialize a *p × p* matrix 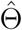, and *K × K* matrices 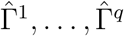.
2. Iterate until convergence:

a. Update Θ by minimizing (4) with respect to Θ, holding 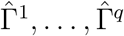 fixed:

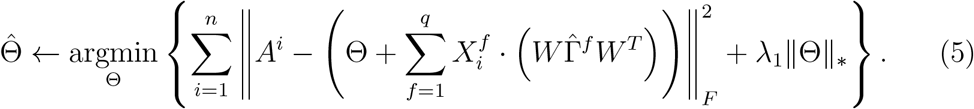
b. For *f* = 1,…, *q*, update Γ^*f*^ by minimizing (4) with respect to Γ^*f*^, holding 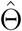 and 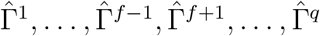 fixed:

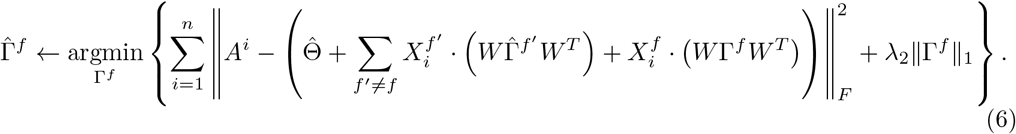

Both (5) and (6) are convex optimization problems, for which closed form solutions are available, as detailed in the following propositions. These propositions make use of the soft-thresholding operator, defined as

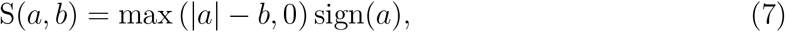

and applied to each element of the matrix.

#### Proposition 1.

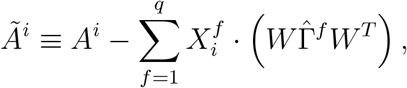

and let *UDV^T^* denote the singular value decomposition of 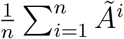: that is, 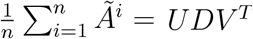, where *U* and *V* are *p × p* matrices, *U^T^U* = *UU_T_* = *V^T^V* = *VV^T^* = *I*, and *D* is a diagonal matrix with non-negative elements on the diagonal. Then, the solution to the optimization problem (5) is

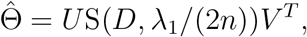

where the soft-thresholding operator defined in (7) is applied element-wise.

Let *p_k_* ≡ |*C_k_*|, the cardinality of the *k*th community; note that 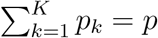.

#### Proposition 2.

Let *W_j_* denote the *j*th row of the matrix *W*. For *f* = 1,…,*q*, define

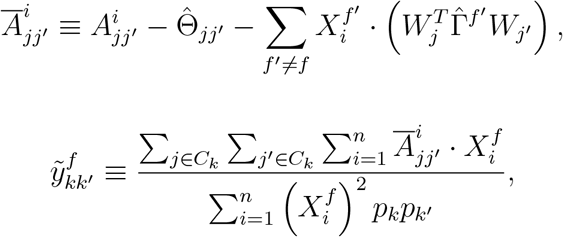

and

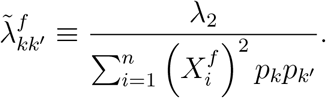

Then, the solution to the optimization problem (6) is of the form

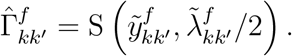

Proofs of Propositions 1 and 2 are provided in the Appendix.

### 2.4. Code Availability

An implementation of the algorithm described above is available in R at bitbucket.org/rshinohara/networkregression.

## 3. Methods

### 3.1. Philadelphia Neurodevelopmental Cohort

Resting-state functional magnetic resonance imaging (rs-fMRI) datasets were acquired as part of the Philadelphia Neurodevelopmental Cohort (PNC), a large community-based study of brain development (Satterthwaite et al. 2014). In total, 1601 participants completed the cross-sectional neuroimaging protocol. Of these participants, 154 were excluded for meeting any of the following criteria: gross radiological abnormalities, history of medical problems that might affect brain function, history of inpatient psychiatric hospitalization, use of psychoactive medications at the time of data acquisition. Of the remaining 1447 participants, 51 were excluded for low quality or incomplete FreeSurfer reconstruction of T1-weighted images. Of the remaining 1396 participants, 381 were excluded for missing rs-fMRI, voxelwise coverage or excessive motion, which is defined as having an average framewise motion of more than 0.20mm and more than 20 frames exhibiting over 0.25mm movement (using calculation from (Jenkinson et al. 2002)). These exclusions produced a final sample consisting of 1015 youth (mean age 15.78, SD = 3.34; 461 males and 554 females).

### 3.2. Imaging Acquisition

Structural and functional imaging data were acquired on a 3T Siemens Tim Trio scanner with a 32-channel head coil (Erlangen, Germany), as previously described (Satterthwaite et al. 2014; Satterthwaite et al. 2016). High-resolution structural images were acquired in order to facilitate alignment of individual subject images into a common space. Structural images were acquired using a magnetization-prepared, rapid-acquisition gradient-echo (MPRAGE) T1-weighted sequence (*T_R_* = 1810ms; *T_E_* = 3.51ms; FoV = 180 × 240mm; resolution 0.9375 × 0.9375 × 1mm). Approximately 6 minutes of task-free functional data were acquired for each subject using a blood oxygen level-dependent (BOLD-weighted) sequence (*T_R_* = 3000ms; *T_E_* = 32ms; FoV = 192 × 192mm; resolution 3mm isotropic; 124 volumes). Prior to scanning, in order to acclimatize subjects to the MRI environment and to help subjects learn to remain still during the actual scanning session, a mock scanning session was conducted using a decommissioned MRI scanner and head coil. Mock scanning was accompanied by acoustic recordings of the noise produced by gradient coils for each scanning pulse sequence. During the mock scanning sessions, feedback regarding head movement was provided using the MoTrack motion tracking system (Psychology Software Tools, Inc, Sharpsburg, PA). To further minimize motion, prior to data acquisition subjects’ heads were stabilized in the head coil using one foam pad over each ear and a third over the top of the head. During the resting-state scan, a fixation cross was displayed as images were acquired. Subjects were instructed to stay awake, keep their eyes open, fixate on the displayed crosshair, and remain still.

### 3.3. Structural Pre-Processing

A study-specific template was generated from a sample of 120 PNC subjects balanced across sex, race, and age using the `buildTemplateParallel` procedure in ANTs (Avants et al. 2011a). Study-specific tissue priors were created using a multi-atlas segmentation procedure (Wang et al. 2013). Next, each subject’s high-resolution structural image was processed using the ANTs Cortical Thickness Pipeline (Tustison et al. 2014). Following bias field correction (Tustison et al. 2010), each structural image was diffeomorphically registered to the study-specific PNC template using the top-performing SyN deformation (Klein et al. 2009). Study-specific tissue priors were used to guide brain extraction and segmentation of the subject’s structural image (Avants et al. 2011b).

### 3.4. Functional Pre-Processing

Task-free functional images were processed using the XCP Engine (Ciric et al. 2017), which was configured to execute a top-performing pipeline for removal of motion-related variance (Ciric et al. 2018). Preprocessing steps included (1) correction for distortions induced by magnetic field inhomogeneities using FSL’s `FUGUE` utility, (2) removal of the 4 initial volumes of each acquisition, (3) realignment of all volumes to a selected reference volume using `mcflirt` (Jenkinson et al. 2002), (4) removal of and interpolation over intensity outliers in each voxel’s time series using AFNI’s `3Ddespike` utility, (5) demeaning and removal of any linear or quadratic trends, and (6) co-registration of functional data to the high-resolution structural image using boundary-based registration (Greve and Fischl 2009). Confounding signals in the data were modelled using a total of 36 parameters, including the 6 framewise estimates of motion, the mean signal extracted from eroded white matter and cerebrospinal fluid compartments, the mean signal extracted from the entire brain, the derivatives of each of these 9 parameters, and the quadratic terms of each of the 9 parameters and their derivatives. Both the BOLD-weighted time series and the artefactual model time series were temporally filtered using a first-order Butterworth filter with a passband between 0.01 and 0.08 Hz (Hallquist, Hwang, and Luna 2013).

### 3.5. Network Construction

We used a common parcellation of cortical and subcortical tissue into 264 regions (Power et al. 2011). The functional connectivity between any pair of brain regions was operationalised as the Pearson correlation coefficient between the mean activation timeseries extracted from those regions. We encoded the pattern of functional connectivity in a formal network model in which nodes represent regions and edges represent functional connections. We assigned each region to one of 13 *a priori* communities (Power et al. 2011) that were delineated using the Infomap algorithm (Rosvall and Bergstrom 2008) and replicated in an independent sample. We excluded 28 nodes that were not sorted into any community, therefore resulting in the final *p* = 236 and *K* = 13. This parcellation was selected for our analysis as it has been previously used for studying individual differences in brain connectivity, including those related to brain development (Gu et al. 2015; Satterthwaite et al. 2012), sex differences (Satterthwaite et al. 2015b), and in-scanner motion (Ciric et al. 2017).

### 3.6. Cross-Validation

We first randomly selected 20% of the total sample (*n* = 1, 015) to serve as the left-out validation set (*n* = 202). We then performed five-fold cross validation on the remaining 80% of the sample (*n* = 813) in order to select the values of the tuning parameters λ_1_ and λ_2_ for MSNR (James et al. 2013). In each fold, the independent variables (*X_n×q_*) were centered to a mean of zero and scaled by each column’s standard deviation. The prediction error used in cross-validation was the *Frobenius* norm of the difference between estimated and true connectivity matrices in the test set, 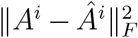. We ensured that the prediction error was independent of the sample size by using the average prediction error over all subjects in the test set.

### 3.7. Permutation Procedure

To estimate the distribution of the prediction error under the null hypothesis of no association between functional connectivity and phenotype, we permuted the rows of the covariate matrix *X_n×q_*. For each permutation, we tuned λ_1_ and λ_2_ using cross-validation, and calculated the prediction error in the left-out validation set. The *p*-value was defined to be the proportion of prediction errors among the 1, 000 permuted datasets that are smaller than the prediction error on the observed data,

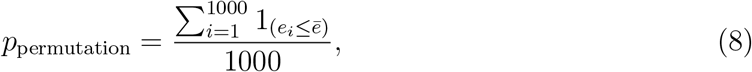

where *e*_1_,…, *e*_1000_ denote the prediction errors on the 1, 000 permuted data sets, and *ē* denotes the prediction error on the original data. Here, 1_(A)_ is an indicator variable that equals 1 if the event *A* holds, and 0 otherwise.

### 3.8. Comparison to Single-Scale Approaches

We compared the performance of MSNR to two of the most commonly used single-scale network regression analyses, namely the individual edge model (Grillon et al. 2013; Lewis et al. 2009) and the community mean model (Yu et al. 2019; Betzel et al. 2014; King et al. 2018; Yan et al. 2019). These two approaches have been commonly used to study connectivity-phenotype relationships (Craddock, Tungaraza, and Milham 2015; Varoquaux and Craddock 2013) and differ primarily in the scale at which the brain network is examined (Fig.3). We describe each model in detail below.

#### Individual edge model

We vectorized the upper triangle of the adjacency matrix *A^i^* for the *i*-th subject, *i* = 1,…, *n*, in order to create a *n* × *p*(*p* – 1)/2 matrix. For each of the *p*(*p* – 1)/2 columns of this matrix, we fit a linear regression in order to model that column using three covariates: age, sex, and in-scanner motion (Fig.3a). Specifically, we built a linear model for each edge using the `mgcv` package in R, with the formula `edge ~ age + sex+ motion` (Wood 2017) (Fig.3b). The model included a penalization on roughness, and we recast the problem as a mixed effect model in order to estimate the penalty parameter via restricted maximum likelihood, or REML (Wood 2011; Wood, Pya, and Safken 2016). We corrected the results for multiple comparisons using the false discovery rate (FDR, *q* < 0.05, (Storey 2002) and reshaped the *p*(*p* – 1)/2 columns to a *p* × *p* matrix in order to visualize significant coefficients. To calculate out-of-sample prediction error, we used linear models fit for all edges. The prediction error was calculated in the same way as in MSNR.

#### Community mean model

Community-based linear models were built with mean *within* - and *between*-community connectivity as the dependent variables. The within-community connectivity is defined as

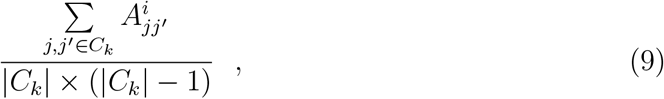

where 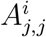 is the weighted edge strength between node *j* and node *j*′, both of which belong to the same community *C_k_*, for the *i*-th subject. The cardinality of the community assignment vector, |*C_k_*|, represents the number of nodes in the *k*-th community. The between-community connectivity is defined as

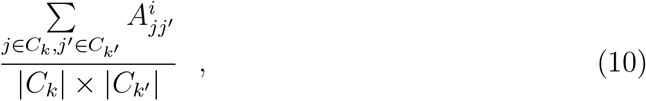

where *C_k_* and *C_k′_* represent two different communities, and |*C_k_*| and 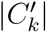 are the number of nodes in each community, respectively.

By applying (9) and (10) to each subject, we created a *n* × [*K*(*K* – 1)/2 + *K*] matrix. For each of the *K*(*K* – 1)/2 + *K* columns of this matrix, we fit a linear model to predict that column using three covariates: age, sex, and in-scanner motion. Similar to the edge-based model, we built a linear model for each edge using the `mgcv` package in R, with the formula `community ~ age + sex+ motion` (Wood 2017) and with roughness penalty estimation performed by REML (Wood 2011; Wood, Pya, and Säfken 2016) (Fig.3b). We corrected the results for multiple comparisons using the false discovery rate (FDR, *q* < 0.05, (Storey 2002)) and reshaped the *K*(*K* – 1)/2 + *K* columns to a *K* × *K* matrix in order to visualize significant coefficients. To calculate out-of-sample prediction error, we used linear models fit for all communitinies. The prediction error was calculated in the same way as in MSNR.

### 3.9. Simulation Study

We used the Brain Connectivity Toolbox (Rubinov and Sporns 2009) to create random modular small-world adjacency matrices of dimension *p* × *p* with specified community assignments (*K* = 4) representing the edge-level information. These adjacency matrices were then used as the ground truth mean connectivity in simulated data, Θ_0_. We also created sparse *K* × *K* matrices 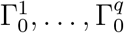, representing ground truth community-level brain-phenotype relationships. We constructed the ground truth adjacency matrix for the *i*th observation as 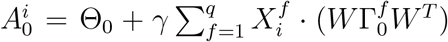, where the elements 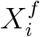 were independently generated from a normal distribution, scaled by a factor of *γ* to represent the effect size. Then, we generated the observed connectivity matrix 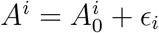 for a noise matrix *ϵ_i_*.

We created synthetic data with varying characteristics, such as different numbers of nodes (*p* ∈ {32, 64,128}), sample sizes (*n* ∈ {50,100,150}), effect sizes (*γ* ∈ {0, 0.1, 0.5,1}), and noise levels (*ϵ* ∈ {0.1, 0.5,1}), for a total of 108 combinations of these parameters. For each combination, we generated three equally-sized sets, for training, testing, and validation.

Tuning parameters λ_1_ and λ_2_ were selected using 1;he training and testing sets, and the out-of-sample prediction error was computed on the validation set.

## 4. Results

### 4.1. MSNR Shows High Accuracy in a Large Developmental Sample

We applied MSNR to data from the Philadelphia Neurodevelopmental Cohort (PNC) (Satterthwaite et al. 2014) in order to delineate known meaningful brain-phenotype relationships. In total, we studied *n* = 1,015 participants aged 8-22 who completed resting state functional neuroimaging as part of the PNC. We constructed functional connectivity matrices from a commonly-used parcellation scheme (*p* = 236 nodes) and community membership assignment (*K* = 13 communities) (Power et al. 2011) (Fig.2A). We first randomly selected 20% of the total sample as the left-out validation set (*n* = 202), with which we assessed the prediction performance of all subsequent models (Fig.2B). The prediction performance was defined as the *Frobenius* norm of the difference between the observed and estimated adjacency matrices in the validation set (Fig.2C). For this proof-of-concept empirical study, we examined the association of functional connectivity with age, sex, and in-scanner motion. On the remaining 80% of the observations, we selected tuning parameters, λ_1_ and λ_2_, through five-fold cross-validation (Fig.2B). We iteratively refined the cross-validation grid (Fig.4A-C) in order to obtain the optimal tuning parameter values. Importantly, no boundary effect was observed during successive grid searches, revealing a smooth convex landscape for the objective (Fig.4D).

**Figure 2:**
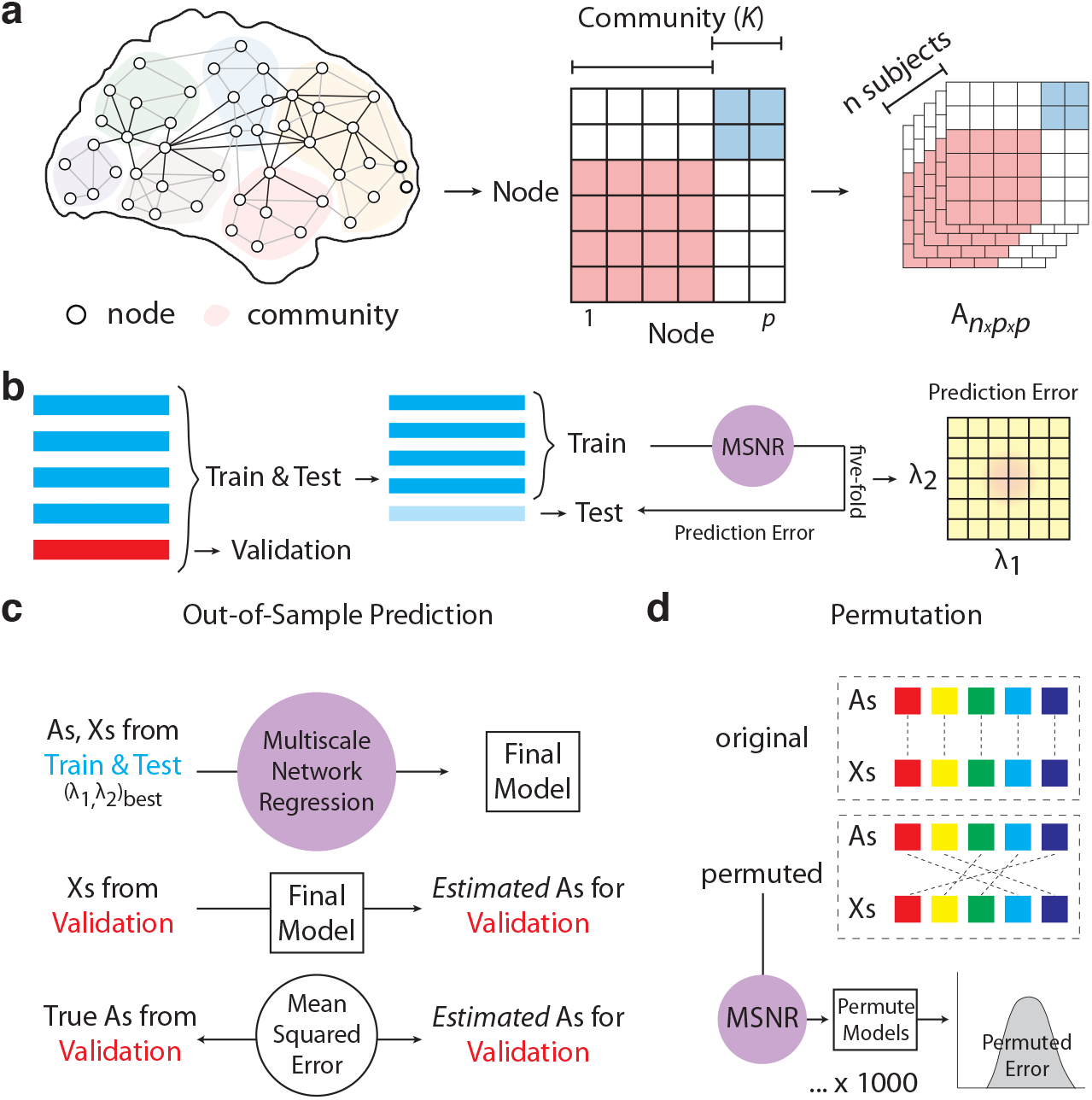
A schematic for MSNR model training and evaluation. **a**) MSNR is designed to study the brain connectivity-phenotype relationship by taking into account both edge- and community-level information. The model takes in a *n × p × p* matrix, where *n* is the number of subjects and where *p* is the number of nodes in each symmetric adjacency matrix. The nodes belong to *K* communities, determined *a priori*. **b**) 20% (*n* = 202) of the total sample (*n* = 1,015) were randomly selected as the left-out validation data. We conducted five-fold cross-validation to select the values of the tuning parameters λ_1_ and λ_2_, which were applied to the nuclear norm penalty on the mean connectivity matrix (Θ) and the *ℓ*_1_ norm of the community-level connectivity-covariate relationship matrices (Γ^1^,…, Γ^*q*^), respectively. **c**) The model was then trained using the tuning parameters determined in **b**) on the 80% (n=813) of the total data not in the left-out validation set. Out-of-sample prediction error was then calculated as the *Frobenius* norm of the difference between the known and estimated connectivity matrices on the validation set. **d**) We also evaluated the final model through a permutation procedure, where we broke the linkage between brain connectivity and covariate data to generate a null distribution of out-of-sample prediction error.

**Figure 3:**
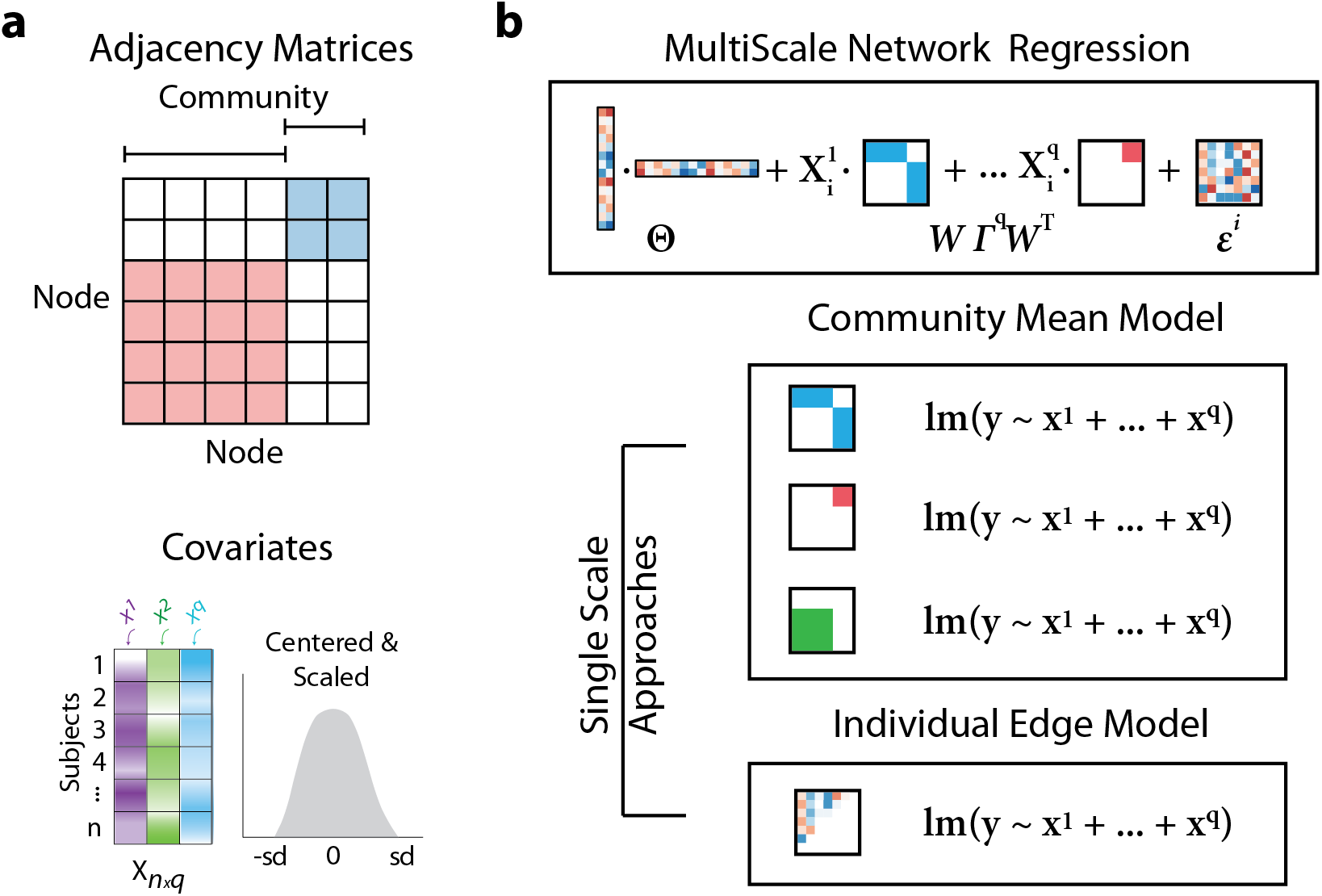
Benchmarking MSNR against common single-scale approaches. **(a)** On the PNC data, we considered prediction of out-of-sample connectivity matrices from age, sex, and in-scanner motion. Specifically, input network data were *n* × *p* × *p* connectivity matrices of n subjects with *p* nodes sorted *a priori* into *K* communities. Additionally, covariate data were a *n* × *q* matrix of *q* measurements, with each column centered with zero mean and scaled by its standard deviation. **(b)** Specifically, we compared MSNR to two common network analysis approaches that only consider informtion present on a single scale. Linear models were fit for each edge or community connectivity for the individual edge and community mean model, respectively.

**Figure 4:**
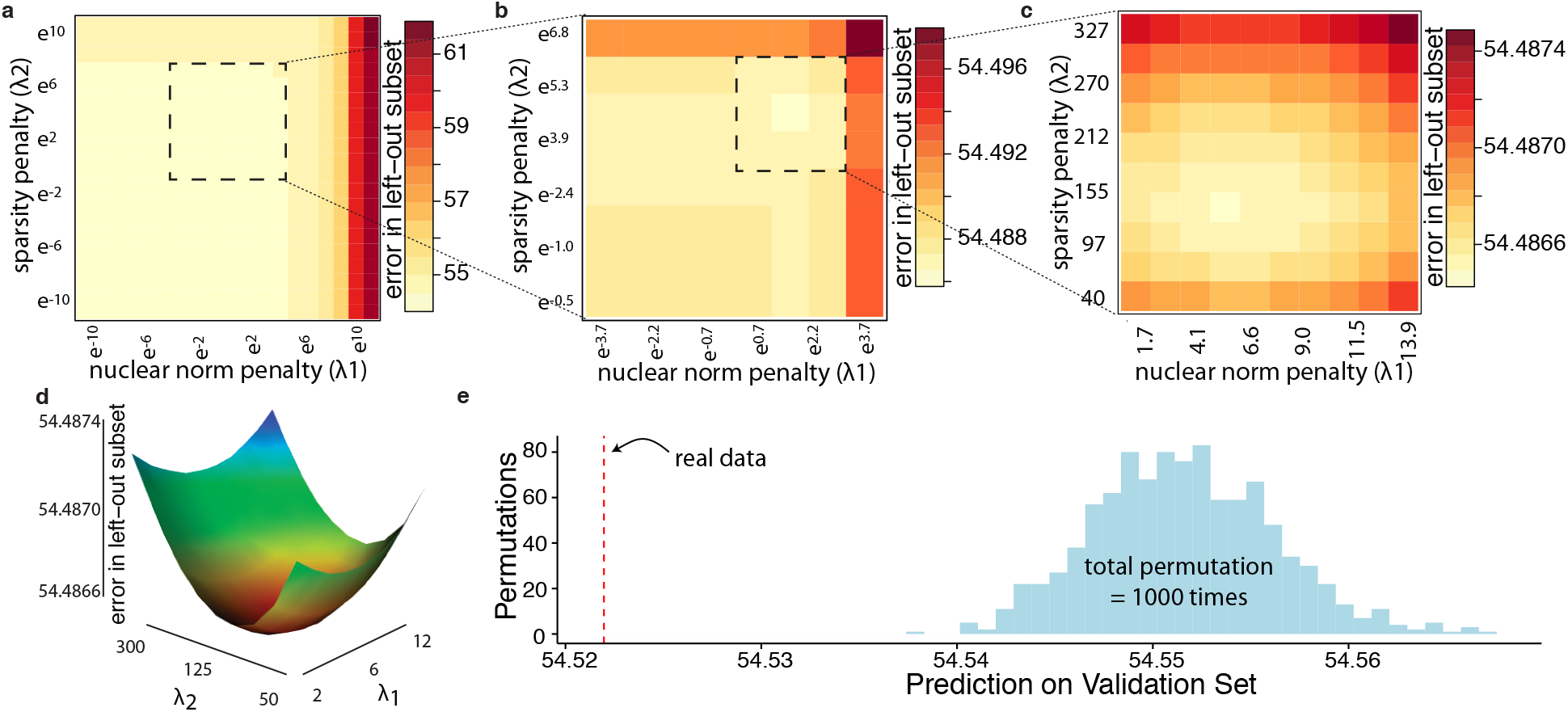
Tuning parameter selection and model evaluation of MSNR in a real-world large neuroimaging dataset. **a)** We used five-fold cross-validation to estimate the test prediction error associated with various values of λi and λ_2_. **b)** After the initial search, we conducted another search on a finer scale, focusing on the range of λ_1_ and λ_2_ indicated by the dashed-line box. **c)** The optimal tuning parameter values were found to be λ_1_ = 5.76 and λ_2_ = 135. No boundary effect was observed in the grid search, revealing a smooth convex landscape for the objective, also visualized in **d)**, with warmer color indicating lower prediction error. **e)** The permutation procedure indicated that MSNR fit to the original data significantly outperformed MSNR fit to permuted data, with a out-of-sample prediction error about six standard deviations below the mean of the null distribution (*p* < 0.001).

We subsequently evaluated the model’s out-of-sample prediction error on the validation set. The prediction error on the unseen data was comparable to the average error in the cross-validation procedure, indicating MSNR did not overfit to the training data (Fig.4E). In addition, to determine the statistical significance of the model, we performed a permutation test to compare the model’s prediction error to the distribution of prediction errors under the null hypothesis of no association between brain networks and the predictors (Fig.4E). Specifically, we permutted the rows of the covariate data matrix 1000 times, which disrupted the linkage between functional connectivity and phenotypes, while preserving the covariance structure of the covariates. For each permutation, we repeated the process of selecting tuning parameter values by cross-validation. The permutation procedure indicated that MSNR fit to the original data significantly outperformed MSNR fit to permuted data, with a out-of-sample prediction error of about six standard deviations below the mean of the null distribution (*p* < 0.001).

### 4.2. MSNR Recapitulates Known Individual Differences in Functional Connectivity

Next, we investigated the connectivity-phenotype relationships that are summarized in the matrices Γ^1^, Γ^2^, and Γ^3^ of the MSNR model. We counted the number of positive and negative coefficients within each estimated matrix; these represent, respectively, positive and negative associations between community pair connectivity with age, sex, and in-scanner motion (Fig.5). Consistent with the previous literature (Satterthwaite et al. 2013; Gu et al. 2015), we found that as age increased, there were more within-, rather than between-community pairs that strengthened connectivity with age (Fig.5A). Conversely, as age increased, there were more between-, rather than within-community pairs that weakened with age. This pattern of results suggests that functional brain networks tend to segregate during normative brain development. Replicating findings from a previous report using mass-univariate analyses (Satterthwaite et al. 2015a), here we observed that stronger within-community connectivity, rather than between-community connectivity, was more representative of functional brain networks in males; whereas stronger between-community connectivity, rather than within-community, was more representative of functional brain networks in females (Fig.5B). Finally, following on prior studies, we evaluated the degree to which the association between in-scanner motion and connectivity varied by inter-node distance, defined as the Euclidean distance between two spherical brain parcellations in the MNI space (Brett, Johnsrude, and Owen 2002) (Fig.5C). As expected, the MSNR coefficients for in-scanner motion in relation to functional connectivity were negatively correlated with the distances between pairs of communities. In other words, when two brain regions were close together, the presence of in-scanner motion was typically associated with an increase in their connectivity. This finding is consistent with prior reports that in-scanner motion in-duces a distance-dependent bias in the estimation of functional connectivity (Satterthwaite et al. 2012; Ciric et al. 2017).

**Figure 5:**
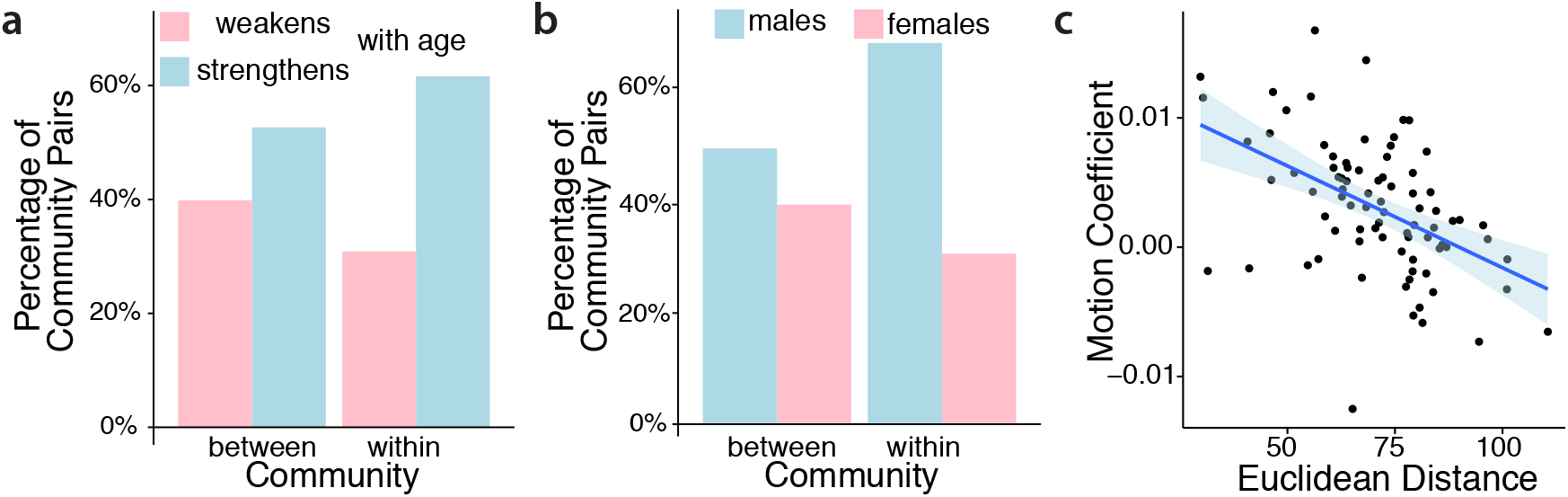
MSNR describes meaningful individual differences in brain connectivity. **a)** More within-community, rather than between-community, connectivity strengthened as the age increased. Conversely, more between-community, rather than within-community, connectivity weakened over age. **b)** Stronger within-community than between-community connectivity was more representative of male functional brain networks, whereas stronger between-community than within-community connectivity was more representative of female functional brain networks. **c)** Coefficient for in-scanner motion was negatively correlated with the average Euclidean distance between communities (*p* < 0.001).

### 4.3. Comparison With Typical Mass-Univariate Single-Scale Strategies

Next, we compared MSNR to common single-scale mass-univariate approaches that make use of linear models at the edge-level or at the community-level (Fig.6). We computed the out-of-sample performance of two single-scale approaches using the left-out validation set. The prediction error of the community-based model on the validation set was poor, whereas that of the edge-based model was similar to MSNR (Fig.6A). Our estimation of prediction error for edge- and community-based models were likely to be overly optimistic, since we used all fitted models for the purpose of out-of-sample prediction. Next, we examined the interpretability of coefficients obtained in each model after applying FDR correction to control for multiple comparisons in single-scale approaches. We found that while the edge-based model and MSNR achieved similar out-of-sample prediction, coefficients estimated in MSNR (Fig.6B) were more interpretable than the coefficients estimated from edge-based models (Fig.6C). The number of coefficients in edge-based models for each covariate exceeded that of MSNR by three orders of magnitude. On the other hand, at the expense of higher prediction error, community-based models exhibited a level of interpretability that was similar to that exhibited by MSNR (Fig.6D).

**Figure 6:**
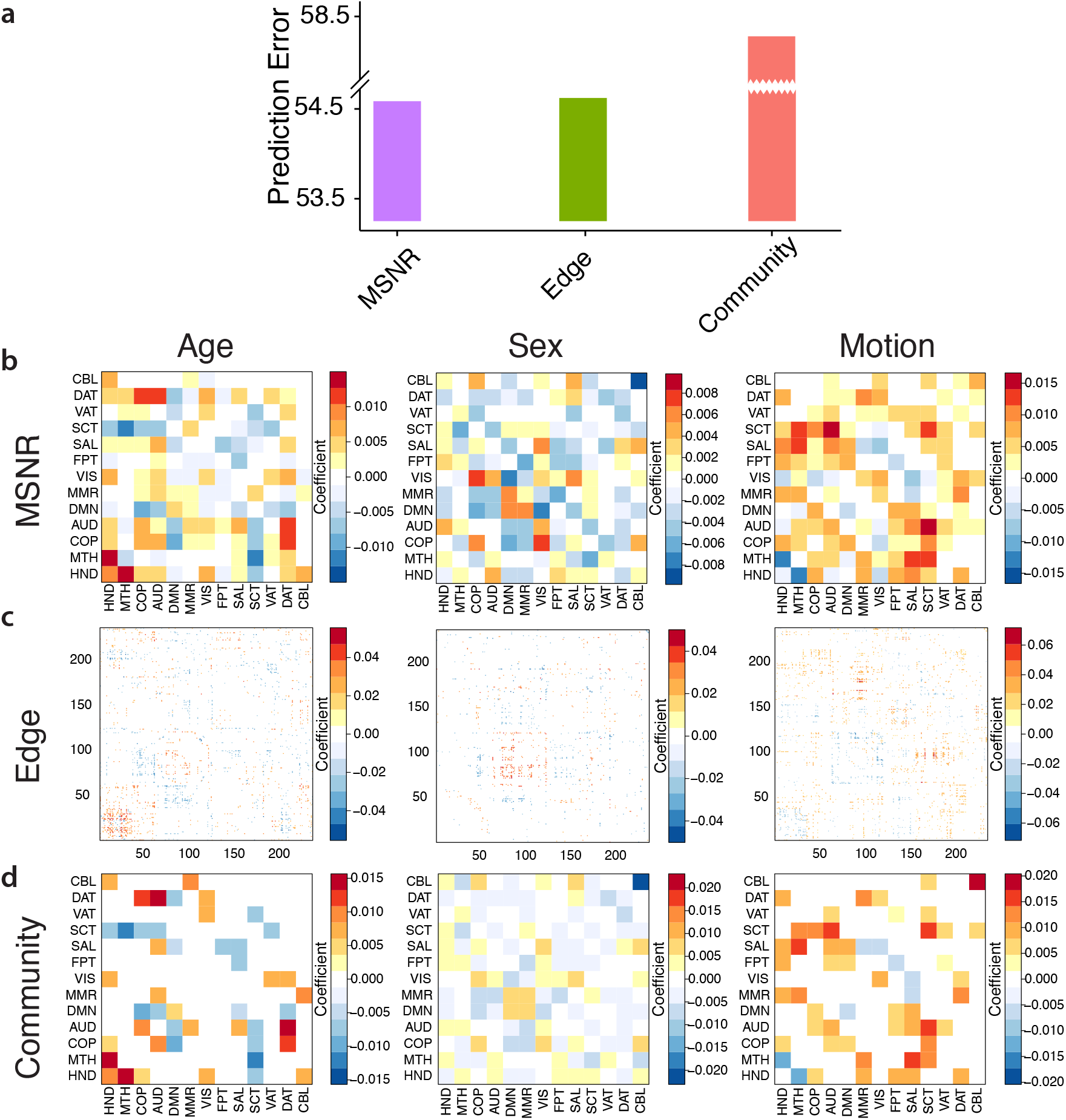
MSNR achieves a balance between out-of-sample prediction performance and model interpretability compared to common single-scale mass-univariate approaches. **a)** We compared out-of-sample prediction performance of MSNR to edge- and community-based single-scale approaches. The community-based approach performed poorly, while the edge-based approach and MSNR had similar out-ofsample prediction error. All models fitted in mass-univariate approaches were used to calculate prediction error. **b)** MSNR coefficients in Γ^1^, Γ^2^, Γ^3^, correspond to age, sex, and in-scanner motion, respectively. Warm colors indicate increased connectivity and cold colors indicate decreased connectivity as the covariate increased. White color indicates zero values. Results from single-scale models were visualized in **c)** for edge-based and in **d)** for community-based approaches. Multiple comparisons were corrected using FDR.

### 4.4. MSNR Behaves As Expected In Simulation Studies

To understand the ability of MSNR to uncover brain-phenotype relationships, we generated adjacency matrices that have a known relationship with randomly-generated covariates (Rubinov and Sporns 2009). Specifically, we simulated adjacency matrices with modular small-world community structures that span a range of number of nodes (*p*), sample sizes (*n*), effect size of the covariates (*γ*), and magnitude of the noise (*ϵ*). We found that MSNR achieved the lowest out-of-sample prediction error when the ratio between the number of subjects and the number of nodes was the largest (*n* = 150, *p* = 32) (Fig.7). In addition, the amount of noise impacted MSNR’s prediction performance in a graded fashion, with a three-fold difference between the lowest noise level (0.1) and the highest noise level (1). In contrast, MSNR was less sensitive to the varying levels of *γ*, which represents the effect size of the community level relationship of the covariates. These results were to be expected, as when the model is well specified in the sense that the data is generated according to the model, the more observations available or the smaller the noise means one can estimate the model parameters more accurately.

**Figure 7:**
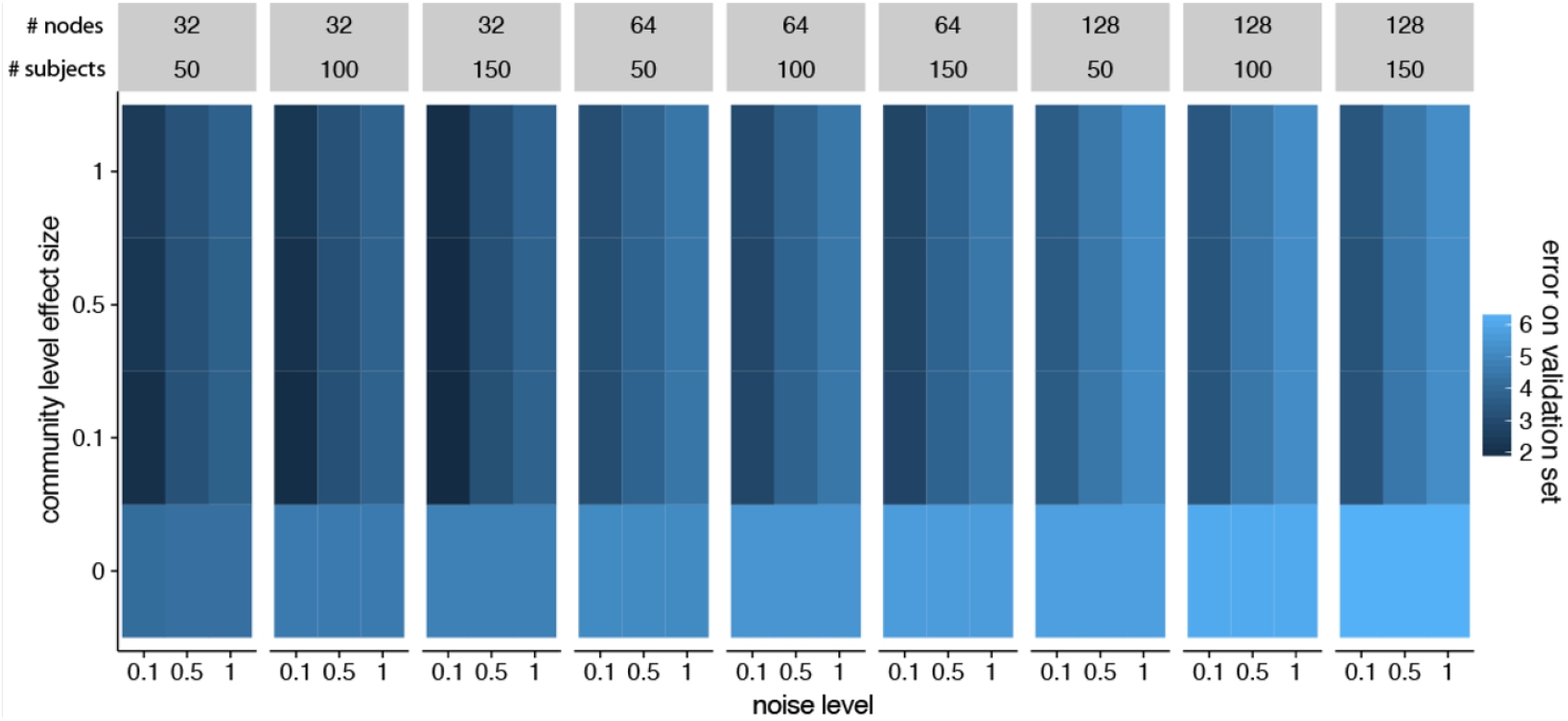
Performance of MSNR in a simulation study. We simulated data with varying numbers of observations (*n*) and nodes (*p*), effect size (*γ*) of Γ^1^ … Γ^*q*^, and noise levels (*ϵ*). As expected, the performance of MSNR improved as the ratio of *n* to *p* increased, and as the signal-to-noise ratio increased. In contrast, MSNR was less sensitive to the varying levels of *γ*, which represents the effect size of the community level relationship of the covariates.

## 5. Discussion

In the past decade, the neuroscience community has begun to complement the study of localized regions of the brain towards studying inter-regional relationships, or connectivity (Bassett and Sporns 2017; Bzdok et al. 2016). The association of network architecture with development and aging throughout the lifespan (Power et al. 2010; Gu et al. 2015; Betzel et al. 2014), cognition (Crossley et al. 2013; Park and Friston 2013; Bressler and Menon 2010), and neuropsychiatric disorders (Yu et al. 2019; Braun et al. 2016; Fornito, Zalesky, and Breakspear 2015; Grillon et al. 2013; Bassett, Xia, and Satterthwaite 2018; Xia et al. 2018; Kernbach et al. 2018) is of profound interest to the burgeoning network neuroscience literature. These brain-phenotype associations can be studied on the scale of individual edges (*micro*-scale), communities (*meso*-scale), or the network as a whole (*macro*-scale), with most existing approaches for analyzing networks, such as mass-univariate analyses, operate on a single scale.

In recent years, interest has centered on multi-scale modeling approaches (Li et al. 2013; Li et al. 2011; Jenatton et al. 2012), which aim to integrate information across homogeneous regions in the brain while still modeling data on finer scales. These methods have mainly focused on the problem of smoothing without prior knowledge of anatomical or functional parcellations of the brain, and have been adapted for both classification (Romberg et al. 2000) and regression (Li et al. 2011) as well as extended to longitudinal settings (Li et al. 2013).

Building upon these recent work, we developed MSNR to study relationships between high-dimensional brain networks and variables of interest. Specifically, our approach modeled the connectivity matrix for each subject by integrating both *micro* - and *meso*-scale network information. By applying a low-rank assumption to the mean connectivity network (Smith et al. 2015; Leonardi et al. 2013; Li et al. 2009) and a sparsity assumption to the community-level network (Xia et al. 2018; Meunier, Lambiotte, and Bullmore 2010; Newman 2006), we substantially decreased the number of parameters and encourage the detection of interpretable brain-phenotype relationships. Leveraging a large neuroimaging dataset of over one thousand youth, we demonstrated that MSNR recapitulated known individual differences in functional connectivity, including those related to development (Satterthwaite et al. 2013; Gu et al. 2015), sex differences (Satterthwaite et al. 2015a), and in-scanner motion (Satterthwaite et al. 2012; Ciric et al. 2017). Notably, compared to common single-scale mass-univariate regression methods, MSNR achieved a balance between prediction performance and model complexity, with improved interpretability. Together, MSNR represents a new method for identifying individual differences in high-dimensional brain networks.

Several limitations of the MSNR approach should be noted. First, the term *scale* does not have a single definition. In fact, as pointed out by (Betzel, Medaglia, and Bassett 2018), scale can represent at least three different entities depending on the context: multi-scale topological structure, multi-scale temporal structure, and multi-scale spatial structure. In MSNR, we only considered multi-scale topological structure. Incorporating additional information from multiple scales beyond those germane to network topology will likely generate more nuanced and richer models for brain networks. Second, while we carefully conducted a permutation test to assess the statistical significance of the entire model, we did not provide an inferential procedure for determining the association between brain networks and each variable of interest. In particular, MSNR makes no claim regarding the statistical significance of the coefficients in the matrices Γ^1^,…, Γ^q^, which describe the community-level relationships with the covariates. Due to the inclusion of penalty terms in the MSNR framework, making such inferential statements is a challenging open problem.

In summary, by explicitly modeling variability both at the edge and community levels, we developed a multi-scale network regression approach that achieved a balance between the trade-off of prediction and model complexity, potentially offering enhanced interpretability. Empirically, we demonstrated its advantages over alternative methods and illustrated its ability to uncover meaningful signals in a large neuroimaging dataset. Approaches such as MSNR have the potential to yield novel insights into brain-behavior relationships that incorporate realistic multi-scale network architecture.

## Acknowledgment

This project was supported in part by DP5OD009145 (D.M.W.), R01NS085211 (R.T.S.), R01NS060910 (R.T.S.), R01MH112847 (R.T.S., T.D.S.), R21MH106799 (D.S.B., T.D.S.), and R01MH113550 (T.D.S., D.S.B.) from the National Institutes of Health, DMS-1252624 (D.M.W.) and DMS-1352060 (Z.M.) from the National Science Foundation, a Blavatnik Fellowship (C.H.X.), a Simons Investigator Award (D.M.W.), a Sloan Fellowship (Z.M.) and a pilot grant from the Center for Biomedical Computing and Analytics at the University of Pennsylvania(R.T.S.). D.B. was funded by BZ2/2-1, BZ2/3-1, BZ2/ 4-1 from the Deutsche Forschungsgemeinschaft, IRTG2150 and Amazon Web Service grants. The PNC was supported by MH089983 and MH089924. D.S.B. also acknowledges support from the MacArthur Foundation, the Alfred P. Sloan Foundation, the ISI Foundation, the Paul Allen Foundation, the Army Research Laboratory, the Office of Naval Research, and the National Science Foundation.

## Appendix

## Proof of Proposition 1.

Given the definition of *Ã_i_*, (5) reduces to the optimization problem

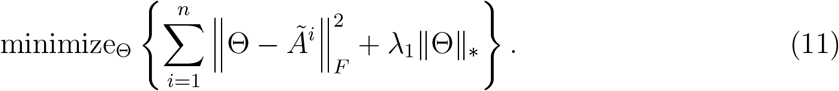

We notice that

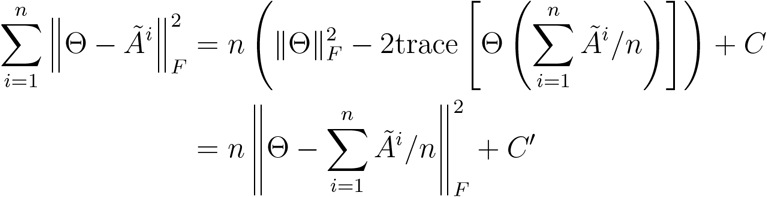

where *C* and *C*′ are not a function of Θ. Therefore, (11) can be re-written as

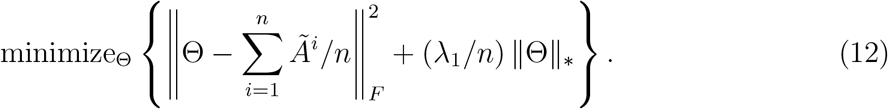

The result follows directly from Lemma 1 of (Mazumder et al. 2010).

## Proof of Proposition 2.

We wish to solve the problem

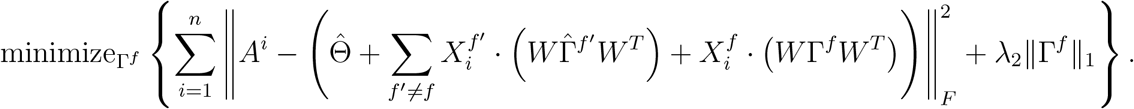

Given the definition of *Ā^i^*, this amounts to solving

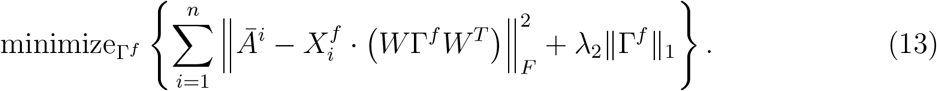

So, for *k* = 1,…, *K* and *k*′ = 1,…, *K*, we must solve the problem

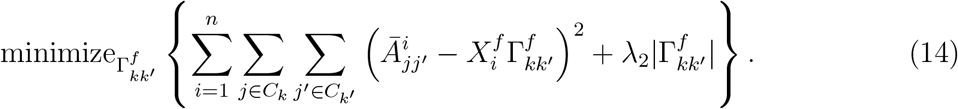

And note that

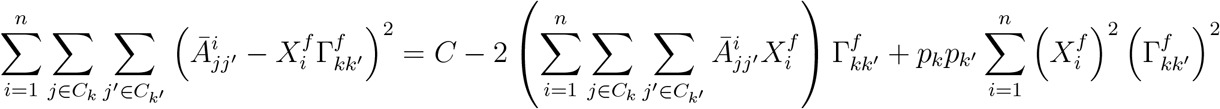

where *C* is not a function of Γ^*f*^. So the problem of interest amounts to minimizing

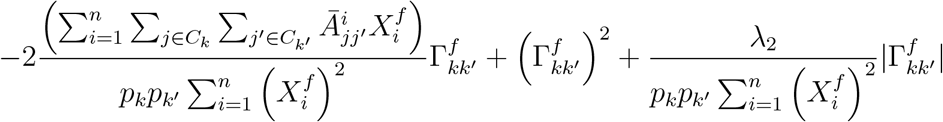

with respect to 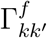. Thus, the minimizer is

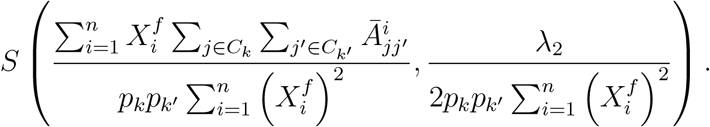

## References

Avants, Brian B., et al. 2011a. “A reproducible evaluation of ANTs similarity metric performance in brain image registration”. NeuroImage 54, no. 3 (): 2033–2044. ISSN: 1053-8119. doi:10.1016/J.NEUR0IMAGE.2010.09.025. https://www.sciencedirect.com/science/article/pii/S1053811910012061.

Avants, Brian B., et al. 2011b. “An Open Source Multivariate Framework for n-Tissue Segmentation with Evaluation on Public Data”. Neuroinformatics 9, no. 4 (): 381–400. ISSN: 1539-2791. doi:10.1007/s12021-011-9109-y. http://link.springer.com/10.1007/s12021-011-9109-y.

Bassett, Danielle S., and Felix Siebenhühner. 2013. “Multiscale Network Organization in the Human Brain”. In Multiscale Analysis and Nonlinear Dynamics, 179–204. Weinheim, Germany: Wiley-VCH Verlag GmbH & Co. KGaA. doi:10.1002/9783527671632.ch07. http://doi.wiley.com/10.1002/9783527671632.ch07.

Bassett, Danielle S., and Olaf Sporns. 2017. “Network neuroscience”. Nature Neuroscience 20 (3): 353–364. doi:10.1038/nn.4502.Network.

Bassett, Danielle S., Cedric Huchuan Xia, and Theodore D. Satterthwaite. 2018. “Understanding the Emergence of Neuropsychiatric Disorders With Network Neuroscience”. Biological Psychiatry: Cognitive Neuroscience and Neuroimaging (). ISSN: 2451-9022. doi:10.1016/J.BPSC.2018.03.015. https://www.sciencedirect.com/science/article/pii/S245190221830079X.

Betzel, Richard F., and Danielle S. Bassett. 2017. “Multi-scale brain networks”. NeuroImage 160 (): 73–83. ISSN: 1053-8119. doi:10.1016/J.NEUROIMAGE.2016.11.006. https://www.sciencedirect.com/science/article/pii/S1053811916306152.

Betzel, Richard F., John D. Medaglia, and Danielle S. Bassett. 2018. “Diversity of meso-scale architecture in human and non-human connectomes”. Nature Communications 9, no. 1 (): 346. ISSN: 2041-1723. doi:10.1038/s41467-017-02681-z. http://www.nature.com/articles/s41467-017-02681-z.

Betzel, Richard F., et al. 2014. “Changes in structural and functional connectivity among resting-state networks across the human lifespan”. NeuroImage 102 (): 345–357. ISSN: 1053-8119. doi:10.1016/J.NEUROIMAGE.2014.07.067. https://www.sciencedirect.com/science/article/pii/S1053811914006508.

Bien, Jacob, and Daniela Witten. 2016. “Penalized estimation in complex models”. Chap. 16 in Handbook of Big Data, 285–299. https://books.google.de/books?hl=en&lr=&id=Kx2VCwAAQBAJ&oi=fnd&pg=PA285&dq=bien+2016+penalized&ots=FKv5nL_kS0&sig=4nbiN10sBeCahzB4Wak_8YlROB8&redir_esc=y#v=onepage&q=bien%202016%20penalized&f=false.

Biswal, Bharat B, et al. 2010. “Toward discovery science of human brain function.” Proceedings of the National Academy of Sciences of the United States of America 107, no. 10 (): 4734–9. ISSN: 1091-6490. doi:10.1073/pnas.0911855107. http://www.ncbi.nlm.nih.gov/pubmed/20176931%20http://www.pubmedcentral.nih.gov/articlerender.fcgi?artid=PMC2842060.

Braun, Urs, et al. 2016. “Dynamic brain network reconfiguration as a potential schizophrenia genetic risk mechanism modulated by NMDA receptor function”. Proceedings of the National Academy of Sciences 113, no. 44 (): 12568–12573. ISSN: 0027-8424. doi:10.1073/pnas.1608819113. http://www.ncbi.nlm.nih.gov/pubmed/27791105%20http://www.pnas.org/lookup/doi/10.1073/pnas.1608819113.

Bressler, Steven L., and Vinod Menon. 2010. “Large-scale brain networks in cognition: emerging methods and principles”. Trends in Cognitive Sciences 14, no. 6 (): 277–290. ISSN: 1364-6613. doi:10.1016/J.TICS.2010.04.004. https://www.sciencedirect.com/science/article/pii/S1364661310000896.

Brett, Matthew, Ingrid S. Johnsrude, and Adrian M. Owen. 2002. “The problem of functional localization in the human brain”. Nature Reviews Neuroscience 3, no. 3 (): 243–249. ISSN: 1471-003X. doi:10.1038/nrn756. http://www.nature.com/articles/nrn756.

Bzdok, Danilo, and John P.A. Ioannidis. 2019. “Exploration, Inference, and Prediction in Neuroscience and Biomedicine”. Trends in Neurosciences 42, no. 4 (): 251–262. ISSN: 01662236. doi:10.1016/j.tins.2019.02.001. http://www.ncbi.nlm.nih.gov/pubmed/30808574%20https://linkinghub.elsevier.com/retrieve/pii/S0166223619300074.

Bzdok, Danilo, and B.T. Thomas Yeo. 2017. “Inference in the age of big data: Future perspectives on neuroscience”. NeuroImage 155 (): 549–564. ISSN: 10538119. doi:10.1016/j.neuroimage.2017.04.061. http://linkinghub.elsevier.com/retrieve/pii/S1053811917303816.

Bzdok, Danilo, et al. 2016. “Formal Models of the Network Co-occurrence Underlying Mental Operations”. Ed. by Danielle S Bassett. PLOS Computational Biology 12, no. 6 (): e1004994. ISSN: 1553-7358. doi:10.1371/journal.pcbi.1004994. https://dx.plos.org/10.1371/journal.pcbi.1004994.

Choi, D. S., P. J. Wolfe, and E. M. Airoldi. 2012. “Stochastic blockmodels with a growing number of classes”. Biometrika 99, no. 2 (): 273–284. ISSN: 0006-3444. doi:10.1093/biomet/asr053. https://academic.oup.com/biomet/article-lookup/doi/10.1093/biomet/asr053.

Ciric, Rastko, et al. 2017. “Benchmarking of participant-level confound regression strategies for the control of motion artifact in studies of functional connectivity”. NeuroImage 154 (March): 174–187. ISSN: 10959572. doi:10.1016/j.neuroimage.2017.03.020.

Ciric, Rastko, et al. 2018. “Mitigating head motion artifact in functional connectivity MRI”. Nature Protocols 13, no. 12 (): 2801–2826. ISSN: 1754-2189. doi:10.1038/s41596-018-0065-y. http://www.nature.com/articles/s41596-018-0065-y.

Craddock, R Cameron, Rosalia L Tungaraza, and Michael P Milham. 2015. “Connectomics and new approaches for analyzing human brain functional connectivity”. GigaScience 4, no. 1 (): 13. ISSN: 2047-217X. doi:10.1186/s13742-015-0045-x. https://academic.oup.com/gigascience/article-lookup/doi/10.1186/s13742-015-0045-x.

Crossley, Nicolas A., et al. 2013. “Cognitive relevance of the community structure of the human brain functional coactivation network”. Proceedings of the National Academy of Sciences 110 (28): 11583–11588. ISSN: 0027-8424. doi:10.1073/pnas.1220826110. http://www.pnas.org/cgi/doi/10.1073/pnas.1220826110.

Durante, Daniele, and David B. Dunson. 2018. “Bayesian Inference and Testing of Group Differences in Brain Networks”. Bayesian Analysis 13, no. 1 (): 29–58. ISSN: 1936-0975. doi:10.1214/16-BA1030. https://projecteuclid.org/euclid.ba/1479179031.

Durante, Daniele, David B. Dunson, and Joshua T. Vogelstein. 2017. “Nonparametric Bayes Modeling of Populations of Networks”. Journal of the American Statistical Association 112, no. 520 (): 1516–1530. ISSN: 0162-1459. doi:10.1080/01621459.2016.1219260. https://www.tandfonline.com/doi/full/10.1080/01621459.2016.1219260.

Fazel, Maryam. 2002. “Matrix rank minimization with applications”. PhD thesis, Stanford University.

Fornito, Alex, Andrew Zalesky, and Michael Breakspear. 2015. “The connectomics of brain disorders”. Nature Reviews Neuroscience 16 (3): 159–172. ISSN: 1471-003X. doi:10.1038/nrn3901. http://dx.doi.org/10.1038/nrn3901.

Fosdick, Bailey K., and Peter D. Hoff. 2015. “Testing and Modeling Dependencies Between a Network and Nodal Attributes”. Journal of the American Statistical Association 110, no. 511 (): 1047–1056. ISSN: 0162-1459. doi:10.1080/01621459.2015.1008697. http://www.tandfonline.com/doi/full/10.1080/01621459.2015.1008697.

Friedman, Jerome, et al. 2007. “Pathwise coordinate optimization”. The Annals of Applied Statistics 1, no. 2 (): 302–332. ISSN: 1932-6157. doi:10.1214/07-AOAS131. http://projecteuclid.org/euclid.aoas/1196438020.

Greve, Douglas N., and Bruce Fischl. 2009. “Accurate and robust brain image alignment using boundary-based registration”. NeuroImage 48, no. 1 (): 63–72. ISSN: 1053-8119. doi:10.1016/J.NEUROIMAGE.2009.06.060. https://www.sciencedirect.com/science/article/pii/S1053811909006752.

Grillon, Marie-Laure, et al. 2013. “Hyperfrontality and hypoconnectivity during refreshing in schizophrenia”. Psychiatry Research: Neuroimaging 211, no. 3 (): 226–233. ISSN: 0925-4927. doi:10.1016/J.PSCYCHRESNS.2012.09.001. https://www.sciencedirect.com/science/article/pii/S0925492712002144.

Gu, Shi, et al. 2015. “Emergence of system roles in normative neurodevelopment”. Proceedings of the National Academy of Sciences 112 (44): 201502829. ISSN: 0027-8424. doi:10.1073/pnas.1502829112. http://www.pnas.org/lookup/doi/10.1073/pnas.1502829112%5Cnhttp://www.pnas.org/content/112/44/13681.abstract.

Hallquist, Michael N., Kai Hwang, and Beatriz Luna. 2013. “The nuisance of nuisance regression: Spectral misspecification in a common approach to resting-state fMRI preprocessing reintroduces noise and obscures functional connectivity”. NeuroImage 82 (): 208–225. ISSN: 1053-8119. doi:10.1016/J.NEUROIMAGE.2013.05.116. https://www.sciencedirect.com/science/article/pii/S1053811913006265.

Hastie, Trevor, Robert Tibshirani, and Jerome Friedman. 2008. The Elements of Statistical Learning. 2nd. Stanford, CA: Springer. https://web.stanford.edu/~hastie/Papers/ESLII.pdf.

Hastie, Trevor, et al. 2015. Statistical Learning with Sparsity. Chapman / Hall/CRC. ISBN: 9781498712170. doi:10.1201/b18401. https://www.taylorfrancis.com/books/9781498712170.

James, Gareth, et al. 2013. An introduction to statistical learning. 8th. Springer. ISBN: 9781461471370. doi:10.1007/978-1-4614-7138-7. www.springer.com.

Jenatton, Rodolphe, et al. 2012. “Multiscale Mining of fMRI Data with Hierarchical Structured Sparsity”. SIAM Journal on Imaging Sciences 5, no. 3 (): 835–856. ISSN: 1936-4954. doi:10.1137/110832380. http://epubs.siam.org/doi/10.1137/110832380.

Jenkinson, Mark, et al. 2002. “Improved Optimization for the Robust and Accurate Linear Registration and Motion Correction of Brain Images”. NeuroImage 17, no. 2 (): 825–841. ISSN: 1053-8119. doi:10.1006/NIMG.2002.1132. https://www.sciencedirect.com/science/article/pii/S1053811902911328.

Jernigan, Terry L., et al. 2016. “The Pediatric Imaging, Neurocognition, and Genetics (PING) Data Repository”. NeuroImage 124, no. Pt B (): 1149–1154. ISSN: 10538119. doi:10.1016/j.neuroimage.2015.04.057. http://www.ncbi.nlm.nih.gov/pubmed/25937488%20http://www.pubmedcentral.nih.gov/articlerender.fcgi?artid=PMC4628902%20http://linkinghub.elsevier.com/retrieve/pii/S1053811915003572.

Kernbach, Julius M., et al. 2018. “Shared endo-phenotypes of default mode dysfunction in attention deficit/hyperactivity disorder and autism spectrum disorder”. Translational Psychiatry 8, no. 1 (): 133. ISSN: 2158-3188. doi:10.1038/s41398-018-0179-6. http://www.nature.com/articles/s41398-018-0179-6.

King, Jace B., et al. 2018. “Evaluation of Differences in Temporal Synchrony Between Brain Regions in Individuals With Autism and Typical Development”. JAMA Network Open 1, no. 7 (): e184777. ISSN: 2574-3805. doi:10.1001/jamanetworkopen.2018.4777. http://jamanetworkopen.jamanetwork.com/article.aspx?doi=10.1001/jamanetworkopen.2018.4777.

Klein, Arno, et al. 2009. “Evaluation of 14 nonlinear deformation algorithms applied to human brain MRI registration”. NeuroImage 46, no. 3 (): 786–802. ISSN: 1053-8119. doi:10.1016/J.NEUROIMAGE.2008.12.037. https://www.sciencedirect.com/science/article/pii/S1053811908012974.

Leonardi, Nora, et al. 2013. “Principal components of functional connectivity: A new approach to study dynamic brain connectivity during rest”. NeuroImage 83 (): 937–950. ISSN: 1053-8119. doi:10.1016/J.NEUROIMAGE.2013.07.019. https://www.sciencedirect.com/science/article/pii/S105381191300774X.

Lewis, Christopher M, et al. 2009. “Learning sculpts the spontaneous activity of the resting human brain.” Proceedings of the National Academy of Sciences of the United States of America 106, no. 41 (): 17558–63. ISSN: 1091-6490. doi:10.1073/pnas.0902455106. http://www.ncbi.nlm.nih.gov/pubmed/19805061%20http://www.pubmedcentral.nih.gov/articlerender.fcgi?artid=PMC2762683.

Li, Kaiming, et al. 2009. “Review of methods for functional brain connectivity detection using fMRI”. Computerized Medical Imaging and Graphics 33, no. 2 (): 131–139. ISSN: 0895-6111. doi:10.1016/J.COMPMEDIMAG.2008.10.011. https://www.sciencedirect.com/science/article/abs/pii/S0895611108001201.

Li, Yimei, et al. 2013. “Multiscale adaptive generalized estimating equations for longitudinal neuroimaging data”. NeuroImage 72 (): 91–105. ISSN: 1053-8119. doi:10.1016/J.NEUROIMAGE.2013.01.034. https://www.sciencedirect.com/science/article/pii/S1053811913000815.

Li, Yimei, et al. 2011. “Multiscale Adaptive Regression Models for Neuroimaging Data.” Journal of the Royal Statistical Society. Series B, Statistical methodology 73, no. 4 (): 559–578. ISSN: 1369-7412. doi:10.1111/j.1467-9868.2010.00767.x. http://www.ncbi.nlm.nih.gov/pubmed/21860598%20http://www.pubmedcentral.nih.gov/articlerender.fcgi?artid=PMC3158617.

Mazumder, Rahul, et al. 2010. Spectral Regularization Algorithms for Learning Large Incomplete Matrices. Tech. rep. https://web.stanford.edu/~hastie/Papers/mazumder10a.pdf.

Meunier, David, Renaud Lambiotte, and Edward T. Bullmore. 2010. “Modular and Hierarchically Modular Organization of Brain Networks”. Frontiers in Neuroscience 4 (): 200. ISSN: 1662-4548. doi:10.3389/fnins.2010.00200. http://journal.frontiersin.org/article/10.3389/fnins.2010.00200/abstract.

Newman, M. E. J. 2006. “Modularity and community structure in networks”. Proceedings of the National Academy of Sciences 103 (23): 8577–8582. ISSN: 0027-8424. doi:10.1073/pnas.0601602103. http://www.pnas.org/cgi/doi/10.1073/pnas.0601602103.

Park, Hae-Jeong, and Karl Friston. 2013. “Structural and functional brain networks: from connections to cognition.” Science (New York, N.Y.) 342, no. 6158 (): 1238411. ISSN: 1095-9203. doi:10.1126/science.1238411. http://www.ncbi.nlm.nih.gov/pubmed/24179229.

Power, Jonathan D., et al. 2010. “The Development of Human Functional Brain Networks”. Neuron 67, no. 5 (): 735–748. ISSN: 0896-6273. doi:10.1016/J.NEURON.2010.08.017. https://www.sciencedirect.com/science/article/pii/S0896627310006276.

Power, JonathanfffdfffdD., et al. 2011. “Functional Network Organization of the Human Brain”. Neuron 72, no. 4 (): 665–678. ISSN: 0896-6273. doi:10.1016/J.NEURON.2011.09.006. https://www.sciencedirect.com/science/article/pii/S0896627311007926.

Raichle, Marcus E. 2015. “The Brain’s Default Mode Network”. Annual Review of Neuroscience 38 (1): 433–447. ISSN: 0147-006X. doi:10.1146/annurev-neuro-071013-014030. arXiv: 0402594v3 [cond-mat]. http://www.annualreviews.org/doi/10.1146/annurev-neuro-071013-014030.

Recht, Benjamin, Maryam Fazel, and Pablo A. Parrilo. 2010. “Guaranteed Minimum-Rank Solutions of Linear Matrix Equations via Nuclear Norm Minimization”. SIAM Review 52, no. 3 (): 471–501. ISSN: 0036-1445. doi:10.1137/070697835. http://epubs.siam.org/doi/10.1137/070697835.

Romberg, J., et al. 2000. “Multiscale classification using complex wavelets and hidden Markov tree models”. In Proceedings 2000 International Conference on Image Processing (Cat. No.00CH37101), 371–374. IEEE. doi:10.1109/ICIP.2000.899396. http://ieeexplore.ieee.org/document/899396/.

Rosvall, Martin, and Carl T Bergstrom. 2008. “Maps of random walks on complex networks reveal community structure”. Proceedings of the National Academy of Sciences of the United States of America 105 (4): 1118–1123. http://www.mapequation.org/assets/publications/RosvallBergstromPNAS2008Full.pdf.

Rubinov, Mikail, and Olaf Sporns. 2009. “Complex network measures of brain connectivity: Uses and interpretations”. NeuroImage 52 (3): 1059–1069. ISSN: 1053-8119. doi:10.1016/j.neuroimage.2009.10.003. http://dx.doi.org/10.1016/j.neuroimage.2009.10.003.

Satterthwaite, Theodore D., et al. 2015a. “Connectome-wide network analysis of youth with Psychosis-Spectrum symptoms.” Molecular psychiatry 20 (February): 1–8. ISSN: 1476-5578. doi:10.1038/mp.2015.66. http://www.ncbi.nlm.nih.gov/pubmed/26033240.

Satterthwaite, Theodore D., et al. 2013. “Heterogeneous impact of motion on fundamental patterns of developmental changes in functional connectivity during youth”. NeuroImage 83 (2013): 45–57. ISSN: 10538119. doi:10.1016/j.neuroimage.2013.06.045. http://dx.doi.org/10.1016/j.neuroimage.2013.06.045.

Satterthwaite, Theodore D., et al. 2012. “Impact of in-scanner head motion on multiple measures of functional connectivity: Relevance for studies of neurodevelopment in youth”. NeuroImage 60 (1): 623–632. ISSN: 10538119. doi:10.1016/j.neuroimage.2011.12.063. http://dx.doi.org/10.1016/j.neuroimage.2011.12.063.

Satterthwaite, Theodore D., et al. 2015b. “Linked Sex Differences in Cognition and Functional Connectivity in Youth”. Cerebral Cortex 25 (9): 2383–2394. ISSN: 14602199. doi:10.1093/cercor/bhu036.

Satterthwaite, Theodore D., et al. 2014. “Neuroimaging of the Philadelphia Neurodevelopmental Cohort”. NeuroImage 86 (2014): 544–553. ISSN: 10959572. doi:10.1016/j.neuroimage.2013.07.064. http://dx.doi.org/10.1016/j.neuroimage.2013.07.064.

Satterthwaite, Theodore D., et al. 2016. “Structural Brain Abnormalities in Youth With Psychosis Spectrum Symptoms”. JAMA Psychiatry 73 (5): 515–524. ISSN: 2168-622X. doi:10.1001/jamapsychiatry.2015.3463. http://archpsyc.jamanetwork.com/article.aspx?doi=10.1001/jamapsychiatry.2015.3463.

Schumann, G, et al. 2010. “The IMAGEN study: reinforcement-related behaviour in normal brain function and psychopathology”. Molecular Psychiatry 15, no. 12 (): 1128–1139. ISSN: 1359-4184. doi:10.1038/mp.2010.4. http://www.ncbi.nlm.nih.gov/pubmed/21102431%20http://www.nature.com/doifinder/10.1038/mp.2010.4.

Smith, Stephen M, et al. 2015. “A positive-negative mode of population covariation links brain connectivity, demographics and behavior”. Nature neuroscience 18 (11): 1565–1567. ISSN: 1097-6256. doi:10.1038/nn.4125. http://dx.doi.org/10.1038/nn.4125.

Sporns, Olaf, and Richard F. Betzel. 2016. “Modular Brain Networks”. Annual Review of Psychology 67 (1): 613–640. ISSN: 0066-4308. doi:10.1146/annurev-psych-122414-033634. http://www.annualreviews.org/doi/10.1146/annurev-psych-122414-033634.

Storey, John D. 2002. “A direct approach to false discovery rates”. Journal of the Royal Statistical Society: Series B (Statistical Methodology) 64, no. 3 (): 479–498. doi:10.1111/1467-9868.00346. http://doi.wiley.com/10.1111/1467-9868.00346.

Tang, Minh, et al. 2017. “A nonparametric two-sample hypothesis testing problem for random graphs”. Bernoulli 23, no. 3 (): 1599–1630. ISSN: 1350-7265. doi:10.3150/15-BEJ789. http://projecteuclid.org/euclid.bj/1489737619.

Tibshirani, Robert. 1996. “Regression Shrinkage and Selection Via the Lasso”. Journal of the Royal Statistical Society: Series B (Methodological) 58, no. 1 (): 267–288. ISSN: 00359246. doi:10.1111/j.2517-6161.1996.tb02080.x. http://doi.wiley.com/10.1111/j.2517-6161.1996.tb02080.x.

Tseng, P. 2001. “Convergence of a Block Coordinate Descent Method for Nondifferentiable Minimization”. Journal of Optimization Theory and Applications 109, no. 3 (): 475–494. ISSN: 0022-3239. doi:10.1023/A:1017501703105. http://link.springer.com/10.1023/A:1017501703105.

Tustison, Nicholas J., et al. 2014. “Large-scale evaluation of ANTs and FreeSurfer cortical thickness measurements”. NeuroImage 99 (): 166–179. ISSN: 1053-8119. doi:10.1016/J.NEUROIMAGE.2014.05.044. https://www.sciencedirect.com/science/article/pii/S1053811914004091.

Tustison, Nicholas J, et al. 2010. “N4ITK: improved N3 bias correction.” IEEE transactions on medical imaging 29, no. 6 (): 1310–20. ISSN: 1558-254X. doi:10.1109/TMI.2010.2046908. http://www.ncbi.nlm.nih.gov/pubmed/20378467%20http://www.pubmedcentral.nih.gov/articlerender.fcgi?artid=PMC3071855.

Van Essen, D.C., et al. 2012. “The Human Connectome Project: A data acquisition perspective”. NeuroImage 62, no. 4 (): 2222–2231. ISSN: 1053-8119. doi:10.1016/J.NEUROIMAGE.2012.02.018. https://www.sciencedirect.com/science/article/pii/S1053811912001954.

Varoquaux, Gafffdfffdl, and R. Cameron Craddock. 2013. “Learning and comparing functional connectomes across subjects”. NeuroImage 80 (): 405–415. ISSN: 1053-8119. doi:10.1016/J.NEUROIMAGE.2013.04.007. https://www.sciencedirect.com/science/article/pii/S1053811913003340.

Wang, Hongzhi, et al. 2013. “Multi-Atlas Segmentation with Joint Label Fusion”. IEEE Transactions on Pattern Analysis and Machine Intelligence 35, no. 3 (): 611–623. ISSN: 0162-8828. doi:10.1109/TPAMI.2012.143. http://ieeexplore.ieee.org/document/6226425/.

Wood, Simon N. 2011. “Fast stable REML and ML estimation of semiparametric GLMs”. Journal of the Royal Statistical Society, Series B (Statistical Methodology) 73 (1): 3–36. ISSN: 13697412. doi:10.1111/j.1467-9868.2010.00749.x.

Wood, Simon N.. 2017. Generalized additive models: an introduction with R. Chapman / Hall/CRC. ISBN: 9781498728331.

Wood, Simon N., Natalya Pya, and Benjamin Safken. 2016. “Smoothing Parameter and Model Selection for General Smooth Models”. Journal of the American Statistical Association 111, no. 516 (): 1548–1563. ISSN: 0162-1459. doi:10.1080/01621459.2016.1180986. https://www.tandfonline.com/doi/full/10.1080/01621459.2016.1180986.

Xia, Cedric Huchuan, et al. 2018. “Linked dimensions of psychopathology and connectivity in functional brain networks”. Nature Communications 9, no. 1 (): 3003. ISSN: 2041-1723. doi:10.1038/s41467-018-05317-y. http://www.nature.com/articles/s41467-018-05317-y.

Yan, Chao-Gan, et al. 2019. “Reduced default mode network functional connectivity in patients with recurrent major depressive disorder.” Proceedings of the National Academy of Sciences of the United States of America (): 201900390. ISSN: 1091-6490. doi:10.1073/pnas.1900390116. http://www.ncbi.nlm.nih.gov/pubmed/30979801.

Young, Stephen J., and Edward R. Scheinerman. 2007. “Random Dot Product Graph Models for Social Networks”. In Algorithms and Models for the Web-Graph, 138–149. Berlin, Heidelberg: Springer Berlin Heidelberg. doi: 10.1007/978-3-540-77004-6{\_}11. http://link.springer.com/10.1007/978-3-540-77004-6_11.

Yu, Meichen, et al. 2019. “Childhood trauma history is linked to abnormal brain connectivity in major depression”. doi:10.1073/pnas.1900801116. www.pnas.org/cgi/doi/10.1073/pnas.1900801116.

Zalesky, Andrew, Alex Fornito, and Edward T. Bullmore. 2010. “Network-based statistic: Identifying differences in brain networks”. NeuroImage 53, no. 4 (): 1197–1207. ISSN: 1053-8119. doi:10.1016/J.NEUROIMAGE.2010.06.041. https://www.sciencedirect.com/science/article/pii/S1053811910008852.

Zapala, Matthew A., and Nicholas J. Schork. 2012. “Statistical Properties of Multivariate Distance Matrix Regression for High-Dimensional Data Analysis”. Frontiers in Genetics 3 (): 190. ISSN: 1664-8021. doi:10.3389/fgene.2012.00190. http://journal.frontiersin.org/article/10.3389/fgene.2012.00190/abstract.

